# Neuroanatomical Risk Factors for Post Traumatic Stress Disorder (PTSD) in Recent Trauma Survivors

**DOI:** 10.1101/721134

**Authors:** Ziv Ben-Zion, Moran Artzi, Dana Niry, Nimrod Jackob Keynan, Yoav Zeevi, Roee Admon, Haggai Sharon, Pinchas Halpern, Israel Liberzon, Arieh Y. Shalev, Talma Hendler

**Affiliations:** Sagol Brain Institute Tel-Aviv, Wohl Institute for Advanced Imaging, Tel Aviv Sourasky Medical Center, Tel-Aviv, Israel; Sagol School of Neuroscience, Tel-Aviv University, Tel-Aviv, Israel; Sackler Faculty of Medicine, Tel-Aviv University, Tel-Aviv, Israel; Department of Radiology, Tel Aviv Sourasky Medical Center, Tel-Aviv, Israel; School of Psychological Sciences, Faculty of Social Sciences, Tel-Aviv University, Tel-Aviv, Israel; Department of Statistics and Operations Research, Tel-Aviv University, Israel; Department of Psychology, University of Haifa, Haifa, Israel; Institute of Pain Medicine, Department of Anesthesiology and Critical Care Medicine, Tel Aviv Sourasky Medical Center, Tel-Aviv, Israel; Pain Management & Neuromodulation Centre, Guy’s & St Thomas’ NHS Foundation Trust, London, UK; Department of Emergency Medicine, Tel Aviv Sourasky Medical Center, Tel-Aviv, Israel; Department of Psychiatry, Texas A&M Health Science Center, Bryan, TX, USA; Department of Psychiatry, NYU Langone Medical Center, New York, NY, USA

**Keywords:** Hippocampus, Cavum Septum Pellucidum, Resilience, Vulnerability, Trauma, Post Traumatic Stress Symptoms

## Abstract

**Background:** Low hippocampal volume could serve as an early risk factor for Post-traumatic Stress Disorder (PTSD) in interaction with other brain anomalies of developmental origin. One such anomaly may well be a presence of large Cavum Septum Pellucidum (CSP), which has been loosely associated with PTSD. Here, we performed a longitudinal prospective study of recent trauma survivors. We hypothesized that at one-month after trauma exposure, the relation between hippocampal volume and PTSD symptom severity will be moderated by CSP volume, and that this early interaction will account for persistent PTSD symptoms at subsequent time-points.

**Methods:** 171 adults (87 females, average age=34.22, range=18-65) admitted to a general hospital’s emergency department following a traumatic event, underwent clinical assessment and structural MRI within one-month after trauma. Follow-up clinical evaluations were conducted at six (n=97) and fourteen (n=78) months after trauma. Hippocampus and CSP volumes were measured automatically by FreeSurfer software and verified manually by a neuroradiologist.

**Results:** At one-month following trauma, CSP volume significantly moderated the relation between hippocampal volume and PTSD severity (p=0.026), and this interaction further predicted symptom severity at fourteen months post-trauma (p=0.018). Specifically, individuals with smaller hippocampus and larger CSP at one-month post-trauma, showed more severe symptoms at one-and fourteen months following trauma exposure.

**Conclusions:** Our study provides evidence for an early neuroanatomical risk factors for PTSD, which could also predict the progression of the disorder in the year following trauma exposure. Such a simple-to-acquire neuroanatomical signature for PTSD could guide early management, as well as long-term monitoring.

**Trial Registration:** Neurobehavioral Moderators of Post-traumatic Disease Trajectories. ClinicalTrials.gov registration: NCT03756545. https://clinicaltrials.gov/ct2/show/NCT03756545

## Introduction

Although accumulating findings point to structural brain abnormalities as potential risk for the development of post-traumatic stress disorder (PTSD), reliable neural measure of the estimated risk (1.3-12%) has yet to be discovered(1–4). Such vulnerability factors could allow accurate diagnosis and therapeutic intervention in the early aftermath of the traumatic event, which has been shown to reduce the likelihood of developing chronic PTSD(5–7).

The most replicated structural abnormality found in PTSD is lower hippocampal volume(8–13), with substantial evidence that this could represent a risk factor for PTSD, including twins studies (14,15). The emerging picture suggests that the hippocampus may have a multifaceted role in PTSD pathogenesis, including the formation and recall of memory traces for contextual information of traumatic events, and providing a representation of safety or danger of the situation(16). While hippocampal volume reduction was observed after trauma exposure(17) and in chronic PTSD(18), increased hippocampal volume was associated with clinical improvement of PTSD symptoms(19). Furthermore, trauma exposure even in the absence of PTSD was shown to be associated with hippocampal volume decrease(17), and further hippocampal volume reduction was seen in chronic PTSD(18). Thus, it is yet to be clearly established whether reduced hippocampal size in PTSD is the result of trauma exposure, represents a risk factor for PTSD, or a combination of both(18,20,21). One possibility is that the hippocampus is not an isolated structural risk factor for PTSD, but that its pathological impact depends on the presence of additional brain anomaly. The accumulating evidence for the involvement of hippocampal abnormalities early on in PTSD may suggest that anatomical anomalies of developmental origin could be of greater relevance.

One such commonly seen anomaly is persistent enlarged Cavum Septum Pellucidum (CSP), known to be related to aberrant brain development(22,23). The CSP, sometimes inaccurately referred to as “fifth ventricle”, is a small cleft filled with cerebrospinal fluid (CSF), located between two thin translucent leaflet membranes, that extends from the anterior part of the corpus callosum to the superior surface of the fornix. During normal development, the fusion of the septi pellucidi occurs within three to six months of age, due to rapid growth of the hippocampal alvei and the corpus callosum(24). However, in some cases the two leaves of the septum pellucidum do not completely fuse, resulting in persistent CSP, which above a certain size, may reflect neurodevelopmental anomaly in midline structures of the brain(22,25) (*see Fig. 1*). Therefore, persistent enlarged CSP in adults may reflect developmental abnormalities of other brain structures bordering the septum pellucidum, such as the hippocampus(24,26). In accordance, schizophrenia patients with an enlarged CSP also showed smaller amygdala and posterior para-hippocampal gyrus volumes (compared to schizophrenia patients without enlarged CSP)(27).

**Figure 1:**
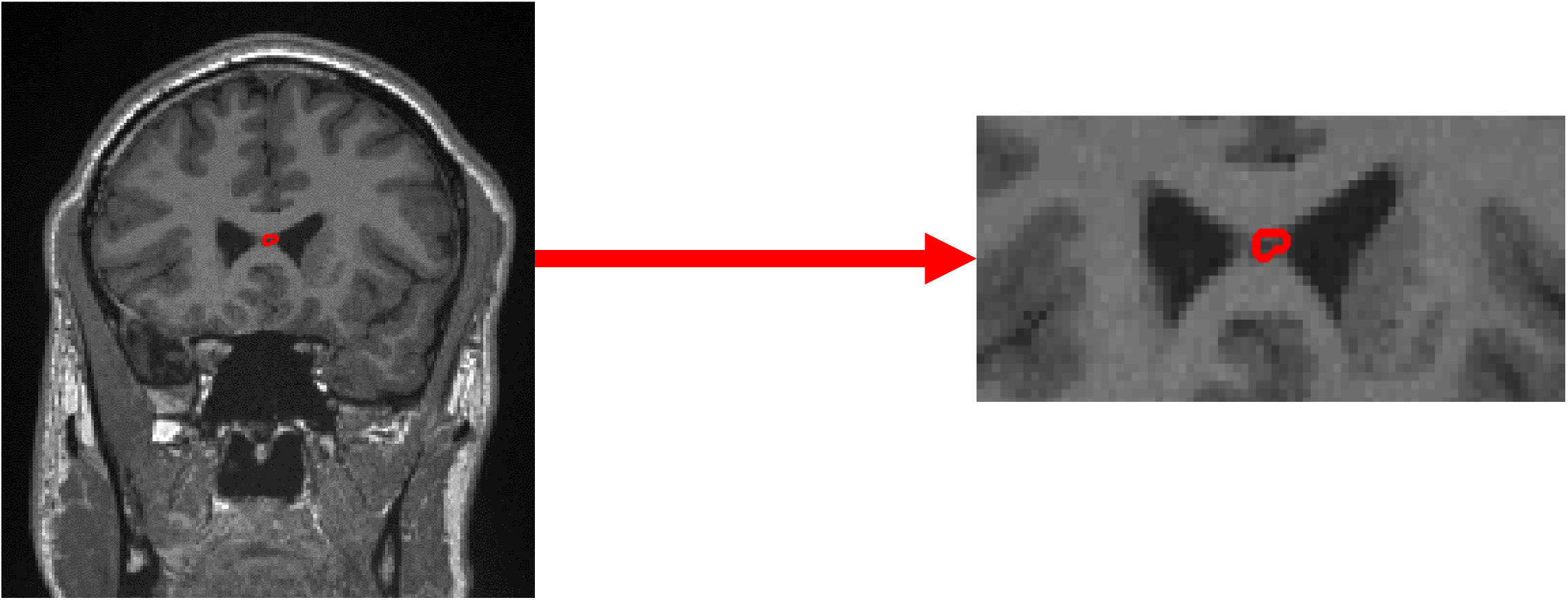
Cavum Septum Pellucidum (CSP). Coronal view of the T_1_-weighted (MPRAGE) image of an example subject. A red line marks the CSP as identified by Freesurfer automatic volumetric segmentation.

One hurdle in pursuing an investigation of the CSP in clinical population, is the uncertainty regarding the prevalence of CSP in healthy adults, stemming from differences in detection method, definition criteria, and homogeneity of the population(23,28–33). Abnormally large CSP was repeatedly linked to schizophrenia(23,28,38,29–32,34–37), bipolar disorders(31,39,40), and others psychopathologies(41,42), when compared to healthy controls. Yet, other studies did not find higher rates of enlarged CSP in schizophrenia(43–45) or other psychiatric disorders(46–49). To date, only two studies have addressed the relationship between the presence of enlarged CSP and PTSD symptomatology. An early study by Myslobodsky et al (1995)(50) has reported increased incidence of CSP (50%) in combat veterans with PTSD, compared with matched normal volunteers (14%), suggesting the CSP might be an antecedent marker for psychopathological vulnerability to stress. More recently, May et al. (2004)(51) reported a greater proportion of enlarged CSP (≥5.6mm) in combat-exposed twins with PTSD and their noncombat-exposed co-twins, suggesting that the presence of an abnormal CSP may serve as a familial vulnerability factor for PTSD.

In the current study we tested the relationship between CSP volume and hippocampus volume in a large cohort of recent trauma survivors, using structural MRI and a within-subject repeated-measures approach. Specifically, we examined hippocampus and CSP volumes within on one-month after trauma, and PTSD symptoms at one-, six-, and fourteen months following trauma exposure. CSP and hippocampal three-dimensional volumes were assessed using automated tools to provide a continuous measure of size of a reliably demarcated region. While for the hippocampus this automated approach has been validated in a number of studies(52–54), for the CSP, as far as we know, no such studies were reported. Therefore, we employed an additional validation procedure by a blinded neuroradiologist for the automated CSP assessment. Our main hypothesis was that the relationship between hippocampal volume and post-traumatic stress symptoms will be moderated by CSP volume, such that individuals with lower hippocampal volume and higher CSP volume would exhibit more severe PTSD symptoms at one-month post-trauma. We used regression models to test this interaction between hippocampal and CSP volumes to predict post-traumatic stress symptoms at one-month post-trauma. Our secondary hypotheses were that this interaction would predict subsequent PTSD symptom severity at six-and fourteen months post-trauma, such that individuals with lower hippocampal volume and higher CSP volume at one-month post-trauma, would exhibit more severe PTSD symptoms at six-and fourteen months post-trauma.

## Methods and Materials

### Participants

The present study is part of a larger on-going project that examines PTSD development in trauma survivors (for full study protocol please see Ben-Zion et al. (2019)(55)). Here we report structural neuroimaging results as pertain to outlined hypotheses obtained from all the participants who completed clinical and neural assessments within one-month following the traumatic incidents (n=171). Out of 171 individuals, to date 97 and 78 currently completed six-and fourteen months post-trauma follow-ups (respectively). Participants were adult survivors of potentially traumatic events, admitted to a Medical Center’s Emergency Room (ER). Individuals were considered for a telephone screening interview if they met the following inclusion criteria: (i) Age 18 – 65 years (ii) Able to read and comprehend native language (iii) Arrived in the ER due to motor-vehicle accidents, bicycle accidents, physical assaults, terrorist attacks, work accidents, large-scale disaster or other potentially traumatic events. To reduce confounds related to concurrent disorders, the exclusion criteria included: (i) survivors with head trauma with coma exceeding a 30 minutes upon ER arrival; (ii) survivors with known medical condition that will interfere with their ability to give informed consent, cooperate with screening and/or treatment; (iii) survivors with claustrophobia, incompatibility for MRI scan, history of substance abuse, current or past psychotic disorder, chronic PTSD; (iv) individuals using psychotropic medication or recreational drugs in the week that precedes the assessment.

### Procedure

A member of the research team identified potentially trauma-exposed individuals using the ER medical records. Within 10–14 days after potential trauma exposure, the identified individuals were contacted by telephone. After verbal consent, risk of PTSD development was assessed using a modified dichotomous version of the PTSD Checklist (PCL) questionnaire, including both DSM-IV and DSM-5 criteria. This interviewer-administered inventory combined both PCL-4 and PCL-5 items, to map directly onto PTSD’s DSM symptom criteria, and was cross-translated to Hebrew. This version showed high internal consistency (0.94) and test-retest reliability (0.82), and was previously used in recent longitudinal studies of recent trauma survivors(55,56). Individuals which met PTSD symptom criteria and did not meet any of the exclusion criteria, received verbal information about the study. They were subsequently invited to participate in both comprehensive clinical assessment and a high-resolution MRI scan, within one-month post-trauma (TP1). Two identical follow-up meetings (including both clinical and neural assessments) were conducted at six-and fourteen months after trauma (TP2 and TP3, respectively). The study was approved by the ethics committee in the local Medical Center (Reference number 0207/14). All participants gave written informed consent in accordance with the Declaration of Helsinki. The study ClinicalTrials.gov registration number is NCT03756545.

### Clinical Assessments

The clinical status of participants was determined by the Clinician-Administered PTSD Scale (CAPS)(57,58), structured clinical interview corresponding to DSM-based PTSD criteria as determined by dimensions of frequency, intensity, and severity of symptoms. An instrument combining both CAPS-4 and CAPS-5 was used, based on DSM-IV and DSM-5 criteria, accordingly. The CAPS contain explicit, behaviorally anchored questions and rating scale descriptors to enhance reliability. It yields a continuous symptom severity score, obtained by summing individual items’ scores (each item ranges from 0-4, with 0 being absent to 4 being extreme/incapacitating). Since there was a very high correlation between CAPS-4 and CAPS-5 total scores across all time-points (*TP1: r=0.962; TP2: r=0.966; TP3: r=0.971; for all: p<0.001*), we report only CAPS-4 total scores as an outcome measure for PTSD symptom severity. CAPS-4 had been extensively used in neuroimaging studies of PTSD to date, and we report it here in order to keep continuity.

### Magnetic Resonance Imaging (MRI)

#### Acquisition

MRI scans were conducted using a Siemens 3T MAGNETOM scanner, located at the Tel Aviv Sourasky Medical Center. In order to assess subcortical and cortical volumes, as well cortical thickness, we used a high resolution sagittal T_1_-weighted magnetization prepared rapid gradient echo (MPRAGE) sequence (TE=2.29 msec, TR=2400 msec, flip angle=8°, FOV =224 mm, Slice Thickness 0.70 mm, voxel size 0.7×0.7×0.7 mm).

#### Analysis

Cortical reconstruction and volumetric segmentation was performed with the FreeSurfer (FS) image analysis suite (59), which is documented and freely available for download online (http://surfer.nmr.mgh.harvard.edu/). Right and left hippocampal and CSP volumes were derived from this process for each subject. The automated hippocampal volumetric measurement by FS was previously shown to have a good agreement with manual hippocampal volumetric assessment, as well as with other automatic methods(53,60,61); however, this was not done for CSP volumetric measurement. In order to validate the automated measurement of the CSP, individuals’ CSP sizes were manually verified by a senior neuroradiologist (D.N.) who was blinded to participants’ clinical symptoms. For each subject, the FS mask of the CSP was evaluated independently according to its correct location and intensity. Based on this blind assessment, participants were divided into two groups: those in which there was agreement between the FS marking and the manual neuroradiologist evaluation (n=133), and those in which there was disagreement between the two (n=28; hence were excluded from the final analysis).

### Statistical Analysis

In order to test whether CSP volume moderated the relationship between bilateral hippocampal volume and PTSD symptoms, moderation analysis including hierarchical multiple regression analysis was conducted using PROCESS macro for SPSS(62,63). In the first step, two independent continuous variables (bilateral hippocampus and CSP volumes at TP1), alongside four covariates (participants’ age, gender, trauma type and intracranial volume (ICV)), were used to predict the dependent continuous variable (PTSD symptom severity as measured by CAPS-4). In the second step of the regression, the centered interaction term between hippocampal volume and CSP volume (i.e. continuous variable) was added to the regression model to test its contribution. When this interaction term significantly predicted PTSD symptoms, it was further illustrated by testing the conditional effects of CSP volume at the different quartiles of hippocampal volume (Q1=25th percentile, Q2=50th percentile=Median, Q3=75th percentile).

In accordance with common norms, the skewed distribution of CSP volume was treated by adding a constant (b=1.25) to the original values of CSP volume, and then a log transformation was performed on these modified values(64). Both bilateral hippocampal volume and CAPS-4 total scores followed a normal distribution, therefore did not require transformations. Furthermore, to reduce the threat of multi-collinearity, both hippocampal and CSP volumes were centered prior to analyses, and an interaction term between these two was created(65).

## Results

A total of 171 participants completed clinical and neuroimaging assessments within one-month following their traumatic incident (TP1). Out of which, 10 individuals were excluded from the analysis due to a missing MPRAGE sequence (n=3), missing clinical data (n=3), or poor-quality structural scan (n=4). The CSP sizes of the remaining 161 participants were manually verified by a senior neuroradiologist (*see MRI Analysis under Methods and Materials*). Based on this blind assessment, for 28 participants (17%) there was a disagreement between the FS marking and the manual neuroradiologist evaluation, hence they were excluded from the analysis. For the remaining 133 participants, there was an agreement between the automatic marking and the manual neuroradiologist evaluation, hence they were included in the final analyses described below. For these n=133 individuals, there was no significant correlation between hippocampal and CSP volumes at TP1 (r=-0.079, p=0.364).

Most of the traumatic events which the participants exhibited were motor-vehicle accidents (n=108, 81%). The other most common types of trauma included bicycle accidents (n=13, 10%) and physical assaults (n=11, 8%). For the follow-up assessments, n=97 and n=78 participants which completed clinical assessments at six-and fourteen months post-trauma (TP2 and TP3; respectively), were included in the final analyses. Out of the 71% individuals diagnosed with PTSD at one-month post-trauma (according to DSM-IV), 31% still had PTSD diagnosis at six months post-trauma, and only 13% were diagnosed with PTSD at fourteen months post-trauma (according to DSM-IV).

No significant differences were found between the 97 individuals which completed TP2 assessments and the 36 which did not, in bilateral hippocampal volume (p=0.747), CSP volume (p=0.491), or ICV (p=0.813). However, significant differences were found between these two groups in initial PTSD symptom severity (p=0.006), such that the 36 individuals which didn’t continue to TP2 assessments showed higher CAPS-4 total scores at TP1 (M=59.03, SD=19.60) compared to the n=97 which returned for TP2 assessments (M=47.05, SD=23.03). Furthermore, no significant differences were found between the 74 individuals which completed TP3 assessments and the 23 which did not, in bilateral hippocampal volume (p=0.220), CSP volume (p=0.990), ICV (p=0.650), or CAPS-4 total scores at TP1 (p=0.637). Finally, no significant differences were found between participants’ age, gender and trauma type across the three time-points (p>0.05 for all). *For full demographic and clinical characteristics of all participants along the three time-points, please see Table 1*.

**Table 1.**
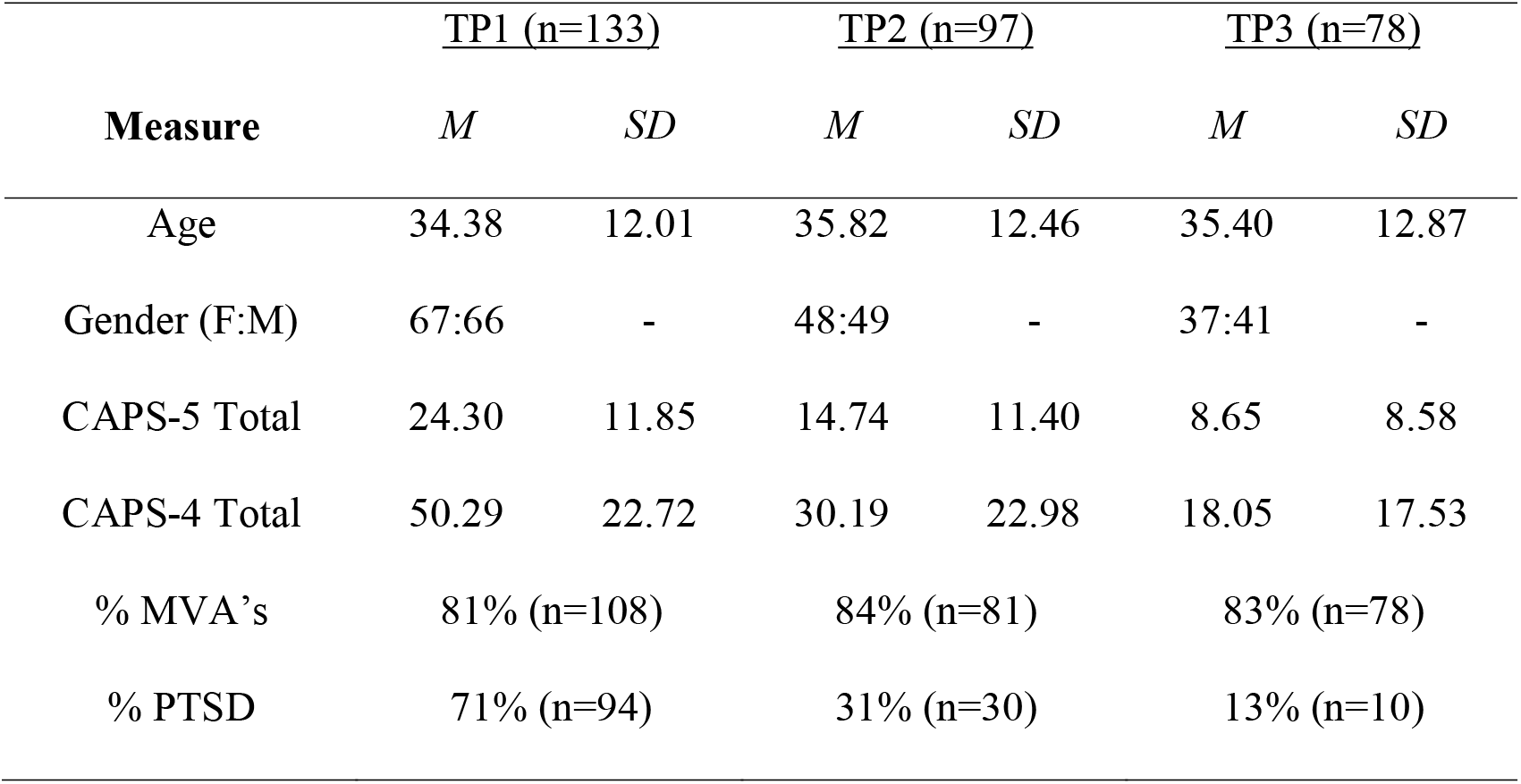
Demographic and clinical characteristics of the participants along the three timepoints. Means (M) and standard deviations (SD) of participants’ age, gender (Female:Male), CAPS-5 and CAPS-4 total scores, at one-, six-and fourteen months following trauma (TP1, TP2 and TP3, respectively). %MVA’s = percentage of motor-vehicle accidents out of all traumatic events; %PTSD = percentage of individuals diagnosed with PTSD (CAPS-4 total score ≥40) out of all participants.

| **Measure** | **<u>TP1 (n=133)</u>** |  | **<u>TP2 (n=97)</u>** |  | **<u>TP3 (n=78)</u>** |  |
| --- | --- | --- | --- | --- | --- | --- |
|  | *M* | *SD* | *M* | *SD* | *M* | *SD* |
| Age | 34.38 | 12.01 | 35.82 | 12.46 | 35.40 | 12.87 |
| Gender (F:M) | 67:66 | - | 48:49 | - | 37:41 | - |
| CAPS-5 Total | 24.30 | 11.85 | 14.74 | 11.40 | 8.65 | 8.58 |
| CAPS-4 Total | 50.29 | 22.72 | 30.19 | 22.98 | 18.05 | 17.53 |
| % MVA's | 81% (n=108) |  | 84% (n=81) |  | 83% (n=78) |  |
| % PTSD | 71% (n=94) |  | 31% (n=30) |  | 13% (n=10) |  |

### Volumetric markers of PTSD symptom severity at one-month after trauma exposure (TP1)

To test our main hypothesis that PTSD symptoms one-month after trauma were associated with multiple volumetric abnormalities, and more specifically whether CSP volume moderates the relationship between hippocampal volume and PTSD severity, a hierarchical multiple regression analysis was conducted (*see details under Statistical Analysis*). Results showed that bilateral hippocampus volume and CSP volume, alongside four covariates (participants’ age, gender, trauma type and ICV), accounted for a significant amount of variance of total scores of CAPS-4 (*R = 0.323, F(6, 126) = 2.439, p = 0.029*). After the interaction term between hippocampal volume and CSP volume was added, the regression model accounted for a significant change in proportion of CAPS-4 total scores (*ΔR^2^= 0.035, ΔF(1,125) = 5.057, p = 0.026*). Consistent with our main hypothesis, a significant interaction (moderation) effect was found between bilateral hippocampus volume and CSP volume in predicting CAPS-4 total scores at one-month post-trauma (*b = −0.0134, t(125) = −2.249, p = 0.026) (See Table 2*). Importantly, neither hippocampal volume nor CSP volume alone predicted CAPS-4 total scores (*p = 0.109; p = 0.183, respectively*).

**Table 2.** Hippocampus volume at TP1 moderates relationship between CSP volume at TP1 and PTSD symptoms at TP1. Regression model of CAPS-4 total scores at TP1 predicted from CSP and hippocampal volumes of 133 participants, with age, gender, trauma type and intracranial volume as covariates. SE=Standard Error.

| **Predictor** | **Beta** | **SE** | **t** | ***p*** |
| --- | --- | --- | --- | --- |
| Bilateral Hippocampus Volume | .0040 | .0025 | 1.6131 | .1092 |
| CSP Volume | 7.9672 | 5.9510 | 1.3388 | .1831 |
| CSP x Hippocampus* | -.0134 | .0060 | -2.2487 | .0263 |
| Age* | -.3374 | .1617 | -2.0867 | .0389 |
| Gender | 2.9068 | 4.8019 | .6054 | .5460 |
| Trauma Type | -.3097 | .9858 | -.3142 | .7539 |
| Intracranial Volume* | .0000 | .0000 | -2.1171 | .0362 |

\* $p < .05$

**Table 3.** Hippocampus volume at TP1 moderates relationship between CSP volume at TP1 and PTSD symptoms at TP3. Regression model of CAPS-4 total scores at TP3 predicted from CSP and hippocampal volumes of 78 participants, with age, gender, trauma type and intracranial volume as covariates. SE=Standard Error.

| **Predictor** | **Beta** | **SE** | **t** | **p** |
| --- | --- | --- | --- | --- |
| Bilateral Hippocampus Volume | -.0001 | .0027 | -.0382 | .9696 |
| CSP Volume | .0297 | 6.1805 | .0048 | .9962 |
| CSP x Hippocampus* | -.0144 | .0059 | -2.4316 | .0176 |
| Age | .0087 | .1583 | .0550 | .9563 |
| Gender | -.2879 | 4.9999 | -.0576 | .9543 |
| Trauma Type | -.2979 | 1.0966 | -.2717 | .7867 |
| Intracranial Volume | .0000 | .0000 | -1.0894 | .2797 |

\* $p < .05$

The above-mentioned interaction was probed by testing the conditional effects of CSP volume at the different quartiles of hippocampal volume (Q1=25th percentile, Q2=50th percentile=Median, Q3=75th percentile) (*see Fig. 2*). At low hippocampal volume (Q1), CSP volume was significantly related to PTSD severity (*CAPS-4: p=0.008*). However, at median and high hippocampal volumes (Q2 and Q3 respectively), the relationship between CSP and hippocampus was not significant (*p>0.15) (see Fig. 2*).

**Figure 2:**
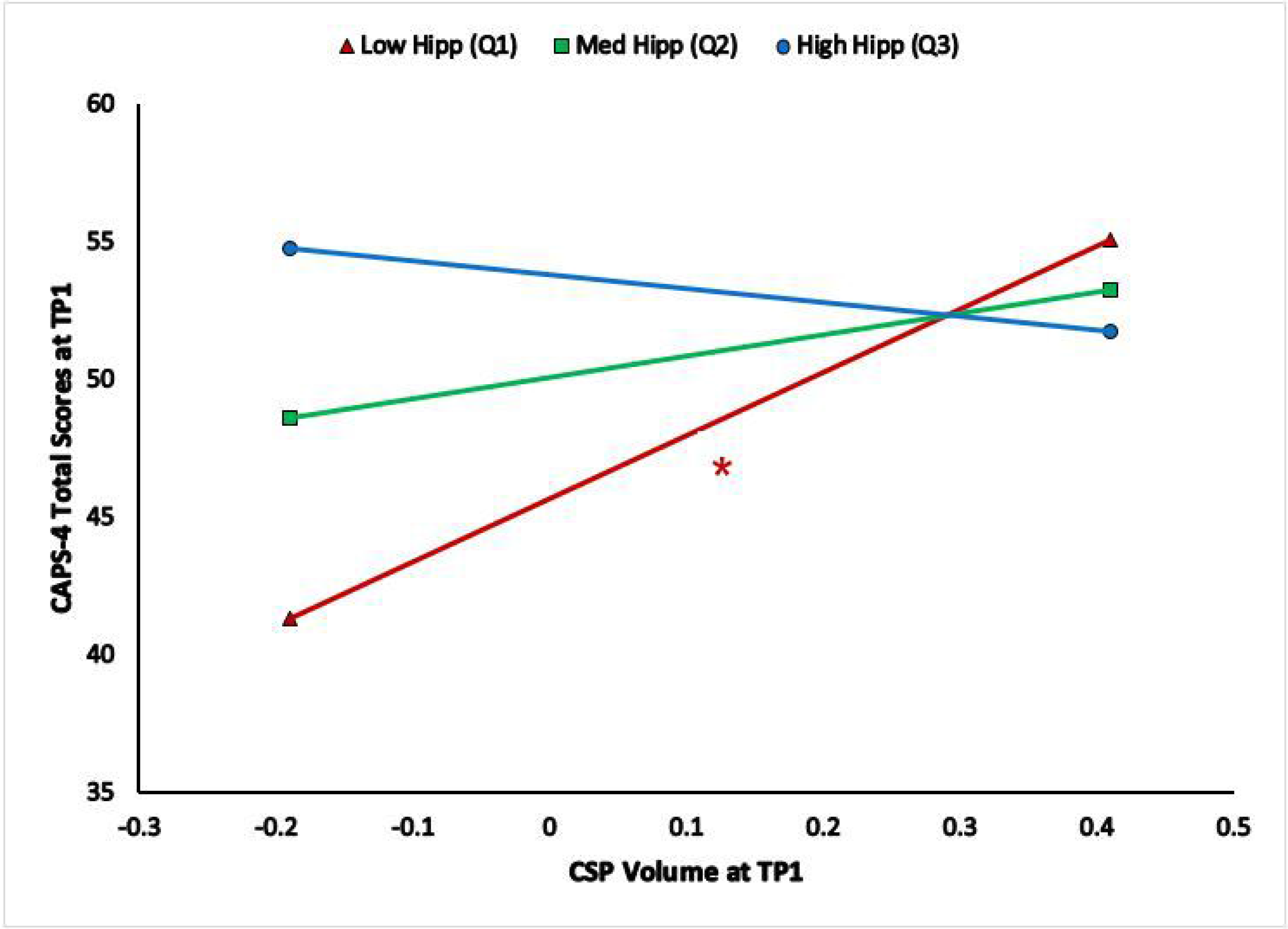
Interaction between hippocampus and CSP volumes at TP1 in predicting TP1 PTSD symptoms. Conditional effects of TP1 CSP volume on TP1 CAPS-4 total scores at different TP1 hippocampal volumes of 133 individuals (Q1=Low Hippocampal Volume in red, Q2=Median Hippocampal Volume in green, Q3=High Hippocampal Volume in blue). Both hippocampal and CSP volumes are centered. Hippocampal volume is presented as a categorical variable with three-levels for illustration purposes, even though it was used as continuous variable in the analyses. *significant at p<0.05

### Volumetric predictors of PTSD symptom severity at follow-up assessments (TP2 and TP3)

To further examine the relation between hippocampal and CSP volumes at TP1 and subsequent PTSD symptoms at TP2 and TP3 (i.e., our secondary hypotheses), two additional hierarchical multiple regression analyses were conducted (one for TP2 and one for TP3).

Focusing on PTSD symptom assessment at six months after trauma (TP2), the hippocampal and CSP volumes at TP1 did not predict PTSD symptom severity at TP2 (hierarchical regression model without the interaction: *R^2^ = 0.080, F(6,90) = 1.304 p = 0.264;* with interaction: *ΔR^2^ = 0.012, ΔF(1,89) = 1.187 p = 0.279*). Hence, contrary to our expectation, there was no significant moderation (interaction) effect between hippocampal and CSP volumes in predicting CAPS-4 total scores at TP2.

With respect to PTSD symptom at fourteen months after trauma (TP3), the hippocampal and CSP volumes at TP1 did not predict PTSD symptom severity at TP3 (*R^2^ = 0.034, F(6,71) = 0.442 p = 0.862);* However, after adding the interaction within a hierarchical regression model, it accounted for a significant proportion of the variance in PTSD symptom severity (*ΔR^2^ = 0.075, ΔF(1,70) = 5.913 p = 0.018*). Consistent with our secondary hypothesis, a significant interaction (moderation) effect was found between bilateral hippocampus and CSP volumes at TP1 in predicting CAPS-4 total scores at TP3 (*b = −0.014, t(70) = −2.432, p = 0.018*) (*See Table 4*). Importantly, neither hippocampal volume nor CSP volume alone predicted CAPS-4 total scores (*p = 0.970; p = 0.996, respectively*).

The above-mentioned interaction was probed by testing the conditional effects of CSP volume at the different quartiles of hippocampal volume (Q1=25th percentile, Q2=50th percentile=Median, Q3=75th percentile) (*see Fig. 3*). At low hippocampal volume (Q1), CSP volume was marginally significantly related to PTSD severity (*p=0.057*). However, at median and high hippocampal volumes (Q2 and Q3 respectively), the relationship between CSP and hippocampus was not significant (*p=0.952 and p=0.276 respectively) (see Fig. 3)*.

**Figure 3:**
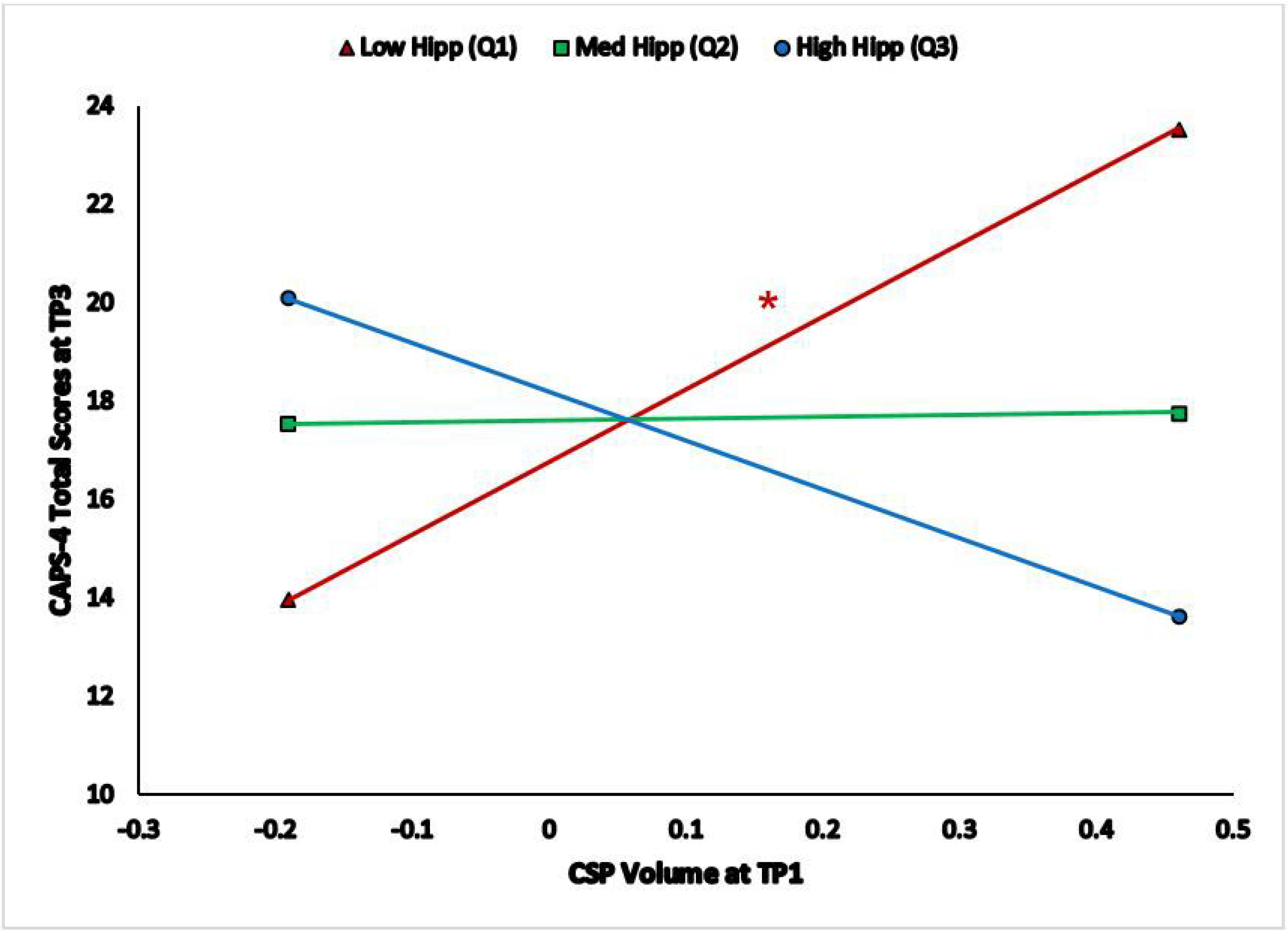
Interaction between hippocampus and CSP volumes at TP1 in predicting TP PTSD symptoms. Conditional effects of TP1 CSP volume on TP3 CAPS-4 total scores at different TP1 hippocampal volumes of 78 individuals (Q1=Low Hippocampal Volume in red, Q2=Median Hippocampal Volume in green, Q3=High Hippocampal Volume in blue). Both hippocampal and CSP volumes are centered. Hippocampal volume is presented as a categorical variable with three-levels for illustration purposes, even though it was used as continuous variable in the analyses. *significant at p<0.10

## Discussion

The current study revealed a moderation effect of CSP volume on the relationship between hippocampal volume and PTSD symptom severity in a population of recent trauma survivors. Specifically, we found that smaller hippocampus volume, together with larger CSP volume, was associated with more severe PTSD symptoms within one-month post-trauma. Furthermore, this relationship at the early aftermath of trauma predicted greater PTSD symptoms at fourteen months following trauma exposure. Together these findings suggest potential neuroanatomical signature of PTSD severity among recent trauma survivors, as well as provide a predictive risk factors for persistent chronicity for the disorder. Our results suggest novel insights regarding the relationship between these two brain structures, recent post-traumatic stress symptoms and predicting chronic course of PTSD. Importantly, their combined (interaction) effect supports a brain development origin for PTSD vulnerability following exposure to potentially traumatic event.

Reduced hippocampal volume is the most consistent finding in structural MRI studies of patients diagnosed chronic PTSD(66–69). Uncertainty exists, however, over the nature and source of smaller hippocampal volumes in PTSD; whether volumetric differences represent the consequence of traumatic exposure, or a pre-existing trait that predisposes people to pathological stress reactions to a traumatic event(70–73). Our results suggest that in the presence of a smaller hippocampus, an abnormally enlarged CSP might serve as a risk factor for developing PTSD following trauma, or vice versa. While a great amount of studies suggests the role of hippocampal volume in PTSD(20), fewer studies have linked abnormal CSP with post-traumatic psychopathology(50,51). We suggest that PTSD vulnerability depends on the interaction between abnormal hippocampal and CSP volumes.

Because an enlarged CSP is considered a neurodevelopmental anomaly, it has been postulated as a potential marker for psychiatric disorders that have neurodevelopmental origins(44). As the postnatal closure of the CSP is dependent on adjacent growing brain structures, a risk for developing different psychopathologies might be associated with a combination of both enlarged CSP and smaller limbic system structures (e.g. hippocampus and amygdala)(74). Here we provide evidence that a combination of enlarged CSP and smaller hippocampal volume might be associated with PTSD symptomatology.

Our study combined early structural MRI indices with longitudinal PTSD clinical measures, enabling to examine the relationship between potential neuroanatomical measures and PTSD symptom severity in the first critical year following trauma. Indeed, we demonstrated that enlarged CSP together with smaller hippocampus measured at one-month following exposure, marked PTSD development and predicted their persistence over 14 months. Nevertheless, the combination of enlarged CSP and smaller hippocampus at one-month post-trauma did not significantly predicted symptom severity at six-month post-trauma, and did not predict change in PTSD symptom severity. This might be explained by the dynamic clinical manifestations during the first critical year following trauma, in which there is a progressive reduction in the severity of PTSD symptoms(75–78). An intermediary point of six-month might be too early to capture the tangible chronic PTSD subtype, whereas 14-months may portray a more stable representation of the chronic disorder as it was shown to predict over 90% of expected PTSD recovery(79,80). Indeed, previous large-scale PTSD symptom trajectories literature reported 17% prevalence of chronic PTSD one year following exposure(5), as was found in our sample (18% out of those who initially suffered from PTSD).

The methodological strengths of the current study derive from the standardized structural MRI measurements obtained in a large cohort-based sample of 171 trauma-exposed individuals with different demographic characteristics (e.g. age, gender). In specific, we applied an automated approach for the volume assessment of CSP yet included only cases that have been also validated by a neuroradiologist (*see MRI Analysis*). This approach, although commonly applied with the hippocampus, goes beyond the current practice regarding CSP measurements. So far, the most commonly applied method has been a subjective classification by a radiologist of small or large CSP(23,28,31,32), dependent on different definitions and criteria, resulting in large variability and inconsistencies among raters. Moreover, this manual classification both time consuming and requires MRI reading expertise. Some researchers adopted more quantitative methods of classification, such as counting the number of slices in which the cavity clearly appears (especially on coronal MRI views), and multiplying it by the slice thickness in order to calculate the anterior-to-posterior length of the cavum(26,27,29,37,43,81). Even with this more quantitate method, there are still conflicting results among the studies that employed such technique(44). Important, such linear measurements only permit a unidimensional representation of the CSP, which could have a complex three-dimensional shape. Volumetric CSP measurement, as employed in this study, may be more meaningful than linear methodologies, since that they provide detailed information about the true size of the structure(26). Our comprehensive approach of combining automated and manual assessments allowed greater confidence in the results and strengthen findings’ generalizability. Having a large sample size allowed to find a sufficient group of individuals with enlarged CSP presence (n=38), thus increasing the statistical power and conclusions.

Although our findings are promising, this work has several limitations. First, as in most PTSD studies, there is a lack of baseline measurement before trauma exposure. However, as structural brain changes usually take time to develop, it is likely to assume that structural abnormalities would reflect pre-disposition factors rather than consequence of trauma exposure. Second, majority of participants suffered from a single trauma, which was mostly related to car accidents. Future work may explore the relationship between these structural brain abnormalities and PTSD symptom severity among varying traumatic events (e.g. terror attacks, sexual or interpersonal violence, continuous traumatic experiences). Lastly, the study included only individuals which experienced early PTSD symptoms, thus are at high-risk for developing chronic PTSD. Future studies should further examine the relation between hippocampus, CSP and PTSD symptomatology in different and larger samples trauma-exposed individuals, in order to increase the validity and replicability of out findings.

This study suggests a promising opportunity of an easy-to-detect individual neuroanatomical abnormalities, large CSP and small hippocampus, that together could serve as distinct neuroanatomical signature for the likelihood of both early and persistent PTSD symptoms following exposure to traumatic events. Such risk factors can be used meaningfully to improve early diagnostic assessment and since it further predicted long term prognosis of the PTSD, it could potentially serve also as a promising monitoring marker for treatment outcome. As structural MRI is becoming more available, we could readily identify individuals who could benefit from early intervention following trauma and follow up their clinical course in an objective manner.

## Acknowledgments

This work was supported by award number R01-MH-103287 from the National Institute of Mental Health (NIMH) given to AS (PI), IL and TH (co-Investigators, subcontractors), and had undergone critical review by the NIMH Adult Psychopathology and Disorders of Aging study section. Nevertheless, the NIMH had no role in the design, execution, data management and data analysis of the current study. Sagol School of Neuroscience at Tel-Aviv University supported ZB fellowships, and Sagol Brain Institute at Tel-Aviv Sourasky Medical Center supported NK and HS fellowships. A preliminary version of this work was previously published in Biorxiv (https://www.biorxiv.org/content/10.1101/721068v1).

## References

1. Shalev A, Liberzon I, Marmar C. Post-Traumatic Stress Disorder. N Engl J Med. 2017;376(25):2459–2469. doi:10.1056/NEJMra1612499

2. Helpman L, Marin MF, Papini S, et al. Neural changes in extinction recall following prolonged exposure treatment for PTSD: A longitudinal fMRI study. NeuroImage Clin. 2016;12:715–723. doi:10.1016/j.nicl.2016.10.007

3. Admon R, Milad MR, Hendler T. A causal model of post-traumatic stress disorder: disentangling predisposed from acquired neural abnormalities. Trends Cogn Sci. 2013;17(7):337–347. doi:10.1016/J.TICS.2013.05.005

4. Swartz JR, Knodt AR, Radtke SR, Hariri AR. A neural biomarker of psychological vulnerability to future life stress. Neuron. 2015;85(3):505–511. doi:10.1016/j.neuron.2014.12.055

5. Galatzer-Levy IR, Ankri Y, Freedman S, et al. Early PTSD Symptom Trajectories: Persistence, Recovery, and Response to Treatment: Results from the Jerusalem Trauma Outreach and Prevention Study (J-TOPS). Felmingham K, ed. PLoS One. 2013;8(8):e70084. doi:10.1371/journal.pone.0070084

6. Shalev AY, Ankri Y, Israeli-Shalev Y, Peleg T, Adessky R, Freedman S. Prevention of posttraumatic stress disorder by early treatment: Results from the Jerusalem trauma outreach and prevention study. Arch Gen Psychiatry. 2012;69(2):166–176. doi:10.1001/archgenpsychiatry.2011.127

7. Shalev AY, Ankri YLE, Peleg T, Israeli-Shalev Y, Freedman S. Barriers to receiving early care for PTSD: results from the Jerusalem trauma outreach and prevention study. Psychiatr Serv. 2011;62(7):765–773. doi:10.1176/appi.ps.62.7.765

8. Pitman RK, Rasmusson AM, Koenen KC, et al. Biological studies of post-traumatic stress disorder. Nat Rev Neurosci. 2012;13(11):769–787. doi:10.1038/nrn3339

9. Kitayama N, Vaccarino V, Kutner M, Weiss P, Bremner JD. Magnetic resonance imaging (MRI) measurement of hippocampal volume in posttraumatic stress disorder: A metaanalysis. J Affect Disord. 2005;88(1):79–86. doi:10.1016/j.jad.2005.05.014

10. Wang Z, Neylan TC, Mueller SG, et al. Magnetic resonance imaging of hippocampal subfields in posttraumatic stress disorder. Arch Gen Psychiatry. 2010;67(3):296–303.

11. Smith ME. Bilateral hippocampal volume reduction in adults with post-traumatic stress disorder: A meta-analysis of structural MRI studies. Hippocampus. 2005;15(6):798–807. doi:10.1002/hipo.20102

12. Karl A, Schaefer M, Malta LS, et al. A meta-analysis of structural brain abnormalities in PTSD. Neurosci Biobehav Rev. 2006;30(7):1004–1031. doi:10.1016/j.amp.2006.02.006

13. Sapolsky RM, Uno H, Rebert CS, Finch CE. Hippocampal damage associated with prolonged glucocorticoid exposure in primates. J Neurosci. 1990;10(9):2897–2902. http://www.ncbi.nlm.nih.gov/pubmed/2398367.

14. Gilbertson MW, Shenton ME, Ciszewski A, et al. Smaller hippocampal volume predicts pathologic vulnerability to psychological trauma. Nat Neurosci. 2002;5(11):1242–1247. doi:10.1038/nn958

15. Gurvits T V., Metzger LJ, Lasko NB, et al. Subtle neurologic compromise as a vulnerability factor for combat-related posttraumatic stress disorder: Results of a twin study. Arch Gen Psychiatry. 2006;63(5):571–576. doi:10.1001/archpsyc.63.5.571

16. Acheson DT, Gresack JE, Risbrough VB. Hippocampal dysfunction effects on context memory: Possible etiology for posttraumatic stress disorder. Neuropharmacology. 2012;62(2):674–685. doi:10.1016/j.neuropharm.2011.04.029

17. Admon R, Leykin D, Lubin G, et al. Stress-induced reduction in hippocampal volume and connectivity with the ventromedial prefrontal cortex are related to maladaptive responses to stressful military service. Hum Brain Mapp. 2013;34(11):2808–2816. doi:10.1002/hbm.22100

18. Woon FL, Sood S, Hedges DW. Hippocampal volume deficits associated with exposure to psychological trauma and posttraumatic stress disorder in adults: A meta-analysis. Prog Neuro-Psychopharmacology Biol Psychiatry. 2010;34(7):1181–1188. doi:10.1016/j.pnpbp.2010.06.016

19. Levy-Gigi E, Szabó C, Kelemen O, Kéri S. Association among clinical response, hippocampal volume, and FKBP5 gene expression in individuals with posttraumatic stress disorder receiving cognitive behavioral therapy. Biol Psychiatry. 2013;74(11):793–800. doi:10.1016/j.biopsych.2013.05.017

20. Pitman RK, Rasmusson AM, Koenen KC, et al. Biological studies of post-traumatic stress disorder. Nat Rev Neurosci. 2012;13(11):769–787. doi:10.1038/nrn3339

21. Bolsinger J, Seifritz E, Kleim B, Manoliu A. Neuroimaging Correlates of Resilience to Traumatic Events—A Comprehensive Review. Front Psychiatry. 2018;9:693. doi:10.3389/fpsyt.2018.00693

22. Bodensteiner JB, Schaefer GB. Wide cavum septum pellucidum: A marker of disturbed brain development. Pediatr Neurol. 1990;6(6):391–394. doi:10.1016/0887-8994(90)90007-N

23. Jurjus GJ, Nasrallah HA, Schwarzkopf SB. Cavum septum pellucidum in schizophrenia, affective disorder and healthy controls: A magnetic resonance imaging study. Psychol Med. 1993;23(2):319–322. doi:10.1017/S0033291700028403

24. Rakic P, Yakovlev PI. Development of the corpus callosum and cavum septi in man. J Comp Neurol. 1968;132(1):45–72. doi:10.1002/cne.901320103

25. Sarwar M. The septum pellucidum: Normal and abnormal. Am J Neuroradiol. 1989;10(5):989–1005. http://www.ncbi.nlm.nih.gov/pubmed/2505543.

26. Crippa JA de S, Zuardi AW, Busatto GF, et al. Cavum septum pellucidum and adhesio interthalamica in schizophrenia: an MRI study. Eur Psychiatry. 2006;21(5):291–299. doi:10.1016/j.eurpsy.2005.09.010

27. Takahashi T, Suzuki M, Hagino H, et al. Prevalence of large cavum septi pellucidi and its relation to the medial temporal lobe structures in schizophrenia spectrum. Prog Neuro Psychopharmacology Biol Psychiatry. 2007;31(6):1235–1241. doi:10.1016/j.pnpbp.2007.04.019

28. Degreef G, Lantos G, Bogerts B, Ashtari M, Lieberman J. Abnormalities of the septum pellucidum on MR scans in first-episode schizophrenic patients. Am J Neuroradiol. 1992;13(3):835–840. http://www.ajnr.org/content/ajnr/13/3/835.full.pdf.

29. Nopoulos P, Swayze V, Flaum M, Ehrhardt JC, Yuh WTC, Andreasen NC. Cavum septi pellucidi in normals and patients with schizophrenia as detected by magnetic resonance imaging. Biol Psychiatry. 1997;41(11):1102–1108. doi:10.1016/S0006-3223(96)00209-0

30. Scott TF, Price TRP, George MS, Brillman J, Rothfus W. Midline cerebral malformations and schizophrenia. J Neuropsychiatry Clin Neurosci. 1993;5(3):287–293. doi:10.1176/jnp.5.3.287

31. Shioiri T, Oshitani Y, Kato T, et al. Prevalence of cavum septum pellucidum detected by MRI in patients with bipolar disorder, major depression and schizophrenia. Psychol Med. 1996;26(02):431. doi:10.1017/S0033291700034838

32. DeLisi LE, Hoff AL, Kushner M, Degreef G. Increased prevalence of cavum septum pellucidum in schizophrenia. Psychiatry Res Neuroimaging. 1993;50(3):193–199. doi:10.1016/0925-4927(93)90030-L

33. Hagino H, Suzuki M, Kurokawa K, et al. Magnetic resonance imaging study of the cavum septi pellucidi in patients with schizophrenia. Am J Psychiatry. 2001;158(10):1717–1719. doi:10.1176/appi.ajp.158.10.1717

34. Fukuzako T, Fukuzako H, Kodama S, Hashiguchi T, Takigawa M. Cavum septum pellucidum in schizophrenia: A magnetic resonance imaging study. Psychiatry Clin Neurosci. 1996;50(3):125–128. doi:10.1111/j.1440-1819.1996.tb01675.x

35. Kwon JS, Shenton ME, Hirayasu Y, et al. MRI study of cavum septi pellucidi in schizophrenia, affective disorder, and schizotypal personality disorder. Am J Psychiatry. 1998;155(4):509–515. doi:10.1176/ajp.155.4.509

36. Nopoulos PC, Giedd JN, Andreasen NC, Rapoport JL. Frequency and severity of enlarged cavum septi pellucidi in childhood-onset schizophrenia. Am J Psychiatry. 1998;155(8):1074–1079. doi:10.1176/ajp.155.8.1074

37. Rajarethinam R, Miedler J, DeQuardo J, et al. Prevalence of cavum septum pellucidum in schizophrenia studied with mri. Schizophr Res. 2001;48(2-3):201–205. doi:10.1016/S0920-9964(00)00110-9

38. Ferrari MCF, Busatto GF, McGuire PK, Crippa JAS. Structural magnetic ressonance imaging in anxiety disorders: An update of research findings. Rev Bras Psiquiatr. 2008;30(3):251–264. doi:10.1109/ICCIS.2013.425

39. Kim MJ, Lyoo IK, Dager SR, et al. The occurrence of cavum septi pellucidi enlargement is increased in bipolar disorder patients. Bipolar Disord. 2007;9(3):274–280. doi:10.1111/j.1399-5618.2007.00442.x

40. Beraldi GH, Prado KS, Amann BL, Radua J, Friedman L, Elkis H. Meta-analyses of cavum septum pellucidum in mood disorders in comparison with healthy controls or schizophrenia. Eur Neuropsychopharmacol. 2018;28(12):1325–1338. doi:10.1016/j.euroneuro.2018.10.001

41. Chun T, Filippi CG, Zimmerman RD, et al. Diffusion changes in the aging human brain. Am JNeuroradiol. 2003;21(6):1078–1083.

42. Nopoulos P, Krie A, Andreasen NC. Enlarged cavum septi pellucidi in patients with schizophrenia: Clinical and cognitive correlates. J Neuropsychiatry Clin Neurosci. 2000;12(3):344–349. doi:10.1176/jnp.12.3.344

43. Flashman LA, Roth RM, Pixley HS, et al. Cavum septum pellucidum in schizophrenia: Clinical and neuropsychological correlates. Psychiatry Res - Neuroimaging. 2007;154(2):147–155. doi:10.1016/j.pscychresns.2006.09.001

44. Trzesniak C, Oliveira IR, Kempton MJ, et al. Are cavum septum pellucidum abnormalities more common in schizophrenia spectrum disorders? A systematic review and meta-analysis. Schizophr Res. 2011;125(1):1–12. doi:10.1016/j.schres.2010.09.016

45. Rajarethinam R, Sohi J, Arfken C, Keshavan MS. No difference in the prevalence of cavum septum pellucidum (CSP) between first-episode schizophrenia patients, offspring of schizophrenia patients and healthy controls. Schizophr Res. 2008;103(1-3):22–25. doi:10.1016/j.schres.2007.11.031

46. Takahashi T, Malhi GS, Wood SJ, et al. Midline brain abnormalities in established bipolar affective disorder. J Affect Disord. 2010;122(3):301–305. doi:10.1016/j.jad.2009.09.003

47. Chon M-W, Choi J-S, Kang D-H, Jung MH, Kwon JS. MRI study of the cavum septum pellucidum in obsessive–compulsive disorder. Eur Arch Psychiatry Clin Neurosci. 2010;260(4):337–343. doi:10.1007/s00406-009-0081-6

48. Takahashi T, Yücel M, Lorenzetti V, et al. Midline brain structures in patients with current and remitted major depression. Prog Neuro-Psychopharmacology Biol Psychiatry. 2009;33(6):1058–1063. doi:10.1016/j.pnpbp.2009.05.020

49. Nopoulos P, Berg S, Castellenos FX, Delgado A, Andreasen NC, Rapoport JL. Developmental Brain Anomalies in Children With Attention-Deficit Hyperactivity Disorder. J Child Neurol. 2000;15(2):102–108. doi:10.1177/088307380001500208

50. Myslobodsky MS, Glicksohn J, Singer J, et al. Changes of brain anatomy in patients with posttraumatic stress disorder: a pilot magnetic resonance imaging study. Psychiatry Res. 1995;58(3):259–264.

51. May FS, Chen QC, Gilbertson MW, Shenton ME, Pitman RK. Cavum septum pellucidum in monozygotic twins discordant for combat exposure: Relationship to posttraumatic stress disorder. Biol Psychiatry. 2004;55(6):656–658. doi:10.1016/j.biopsych.2003.09.018

52. Morey RA, Petty CM, Xu Y, et al. A comparison of automated segmentation and manual tracing for quantifying hippocampal and amygdala volumes. Neuroimage. 2009;45(3):855–866. doi:10.1016/J.NEUROIMAGE.2008.12.033

53. Tae WS, Kim SS, Lee KU, Nam EC, Kim KW. Validation of hippocampal volumes measured using a manual method and two automated methods (FreeSurfer and IBASPM) in chronic major depressive disorder. Neuroradiology. 2008;50(7):569–581. doi:10.1007/s00234-008-0383-9

54. Mulder ER, de Jong RA, Knol DL, et al. Hippocampal volume change measurement: Quantitative assessment of the reproducibility of expert manual outlining and the automated methods FreeSurfer and FIRST. Neuroimage. 2014;92:169–181. doi:10.1016/j.neuroimage.2014.01.058

55. Ben-Zion Z, Fine NB, Keynan NJ, et al. Neurobehavioral Moderators of Post-Traumatic Stress Disorder Trajectories: Prospective MRI Study of Recent Trauma Survivors. Eur J Psychotraumatol. 2019. doi:https://doi.org/10.1080/20008198.2019.1683941

56. Fine NB, Achituv M, Etkin A, Merin O, Shalev AY. Evaluación de la reparación cognitivo-afectiva basada en la web en sobrevivientes recientes de trauma: Fundamentos del estudio y protocolo. Eur J Psychotraumatol. 2018;9(1):1442602. doi:10.1080/20008198.2018.1442602

57. Blake DD, Weathers FW, Nagy LM, et al. The development of a Clinician-Administered PTSD Scale. J Trauma Stress. 1995;8(1):75–90. doi:10.1007/BF02105408

58. Weathers FW, Bovin MJ, Lee DJ, et al. The Clinician-Administered PTSD Scale for DSM–5 (CAPS-5): Development and initial psychometric evaluation in military veterans. Psychol Assess. 2018;30(3):383–395. doi:10.1037/pas0000486

59. Fischl B. FreeSurfer. Neuroimage. 2012;62(2):774–781.

60. Sánchez-Benavides G, Gómez-Ansón B, Sainz A, Vives Y, Delfino M, Peña-Casanova J. Manual validation of FreeSurfer’s automated hippocampal segmentation in normal aging, mild cognitive impairment, and Alzheimer Disease subjects. Psychiatry Res - Neuroimaging. 2010;181(3):219–225. doi:10.1016/j.pscychresns.2009.10.011

61. Wenger E, Mårtensson J, Noack H, et al. Comparing manual and automatic segmentation of hippocampal volumes: Reliability and validity issues in younger and older brains. Hum Brain Mapp. 2014;35(8):4236–4248. doi:10.1002/hbm.22473

62. Preacher KJ, Hayes AF. Asymptotic and resampling strategies for assessing and comparing indirect effects in multiple mediator models. Behav Res Methods. 2008;40(3):879–891. doi:10.3758/BRM.40.3.879

63. Hayes AF. Introduction to Mediation, Moderation and Conditional Process Analysis. Vol 53.; 2013. doi:10.5539/ass.v11n9p207

64. Wang H, Lu N, Chen T, He H, Lu Y, Tu XM. Changyong FENG Log-transformation and its implications for data analysis •Biostatistics in psychiatry (20)•. Shanghai Arch Psychiatry, 2014, Vol 26, No 2. 2014;26(2):105–109. doi:10.3969/j.issn.1002-0829.2014.02.009

65. Aiken LS, West SG, Reno RR. Multiple Regression: Testing and Interpreting Interactions. Vol 45.; 1991. doi:10.2307/2583960

66. Douglas J, Tammy M, Richard A. MRI-based measurement of hippocampal volume in patients with combat-related p … ncbi.nlm.nih.govl 995,;i.

67. Bremner JD, Randall P, Vermetten E, et al. Magnetic resonance imaging-based measurement of hippocampal volume in posttraumatic stress disorder related to childhood physical and sexual abuse - A preliminary report. Biol Psychiatry. 1997;41(1):23–32. doi:10.1016/S0006-3223(96)00162-X

68. Gurvits T, Shenton M, Hokama H, Ohta H, Lasko N, Pitman R. Magnetic resonance imagine study of hippocampal volume in chronic, combat-related PTSD. Biol Psychiatry. 1996;40:1091–1099. http://www.biologicalpsychiatryjoumal.com/article/S0006-3223(96)00229-6/abstract.

69. Stein MB, Koverola C, Hanna C, Torchia MG, McClarty B. Hippocampal volume in women victimized by childhood sexual abuse. Psychological Medicine, 27, 951-959. Psychol Med. 1997;27:951–959.

70. Yehuda R. Are glucocortoids responsible for putative hippocampal damage in PTSD? How and when to decide. Hippocampus. 2001;11(2):85–89. doi:10.1002/hipo.1025

71. McEwen BS. Commentary on PTSD discussion. Hippocampus. 2001;11(2):82–84. doi:10.1002/hipo.1024

72. Douglas Bremner J. Hypotheses and controversies related to effects of stress on the hippocampus: An argument for stress-induced damage to the hippocampus in patients with posttraumatic stress disorder. Hippocampus. 2001;11(2):75–81. doi:10.1002/hipo.1023

73. Pitman RK. Hippocampal diminution in PTSD: More (or Less?) Than meets the eye. Hippocampus. 2001;11(2):73–74. doi:10.1002/hipo.1022

74. Dremmen MHG, Bouhuis RH, Blanken LME, et al. Cavum septum pellucidum in the general pediatric population and its relation to surrounding brain structure volumes, cognitive function, and emotional or behavioral problems. Am J Neuroradiol. 2019;40(2):340–346. doi:10.3174/ajnr.A5939

75. Yehuda R, McFarlane AC, Shalev AY. Predicting the development of posttraumatic stress disorder from the acute response to a traumatic event. Biol Psychiatry. 1998;44(12):1305–1313.

76. Koren D, Arnon I, Klein E. Long term course of chronic posttraumatic stress disorder in traffic accident victims: A three-year prospective follow-up study. Behav Res Ther. 2001;39(12):1449–1458. doi:10.1016/S0005-7967(01)00025-0

77. Hepp U, Moergeli H, Buchi S, et al. Post-traumatic stress disorder in serious accidental injury: 3-Year follow-up study. Br J Psychiatry. 2008;192(5):376–383. doi:10.1192/bjp.bp.106.030569

78. Perkonigg A, Pfister H, Stein MB, et al. Longitudinal course of posttraumatic stress disorder and posttraumatic stress disorder symptoms in a community sample of adolescents and young adults. Am J Psychiatry. 2005;162(7):1320–1327. doi:10.1176/appi.ajp.162.7.1320

79. Kessler RC, Sonnega A, Bromet E, Hughes M, Nelson CB. Posttraumatic Stress Disorder in the National Comorbidity Survey. Arch Gen Psychiatry. 1995;52(12):1048–1060. doi:10.1001/archpsyc.1995.03950240066012

80. Shalev AY, Freedman S. PTSD Following Terrorist Attacks: A Prospective Evaluation. Am J Psychiatry. 2005;162(6):1188–1191. doi:10.1176/appi.ajp.162.6.1188

81. Takahashi T, Yung AR, Yücel M, et al. Prevalence of large cavum septi pellucidi in ultra high-risk individuals and patients with psychotic disorders. Schizophr Res. 2008;105(1-3):236–244. doi:10.1016/j.schres.2008.06.021

